# Defects in AMPAR trafficking and microglia activation underlie socio-cognitive deficits associated to decreased expression of Phosphodiesterase 2A

**DOI:** 10.1101/2023.11.06.565257

**Authors:** Sébastien Delhaye, Marielle Jarjat, Asma Boulksibat, Clara Sanchez, Alessandra Tempio, Andrei Turtoi, Mauro Giorgi, Sandra Lacas-Gervais, Gabriele Baj, Carole Rovere, Viviana Trezza, Manuela Pellegrini, Thomas Maurin, Enzo Lalli, Barbara Bardoni

## Abstract

Phosphodiesterase 2A (PDE2A) is an enzyme involved in the homeostasis of cAMP and cGMP and is the most highly expressed PDE in human brain regions critical for socio-cognitive behavior. In cortex and hippocampus, PDE2A expression level is upregulated in *Fmr1*-KO mice, a model of the Fragile X Syndrome (FXS), the most common form of inherited intellectual disability (ID) and autism spectrum disorder (ASD). Indeed, PDE2A translation is negatively modulated by FMRP, whose functional absence causes FXS. While the pharmacological inhibition of PDE2A has been associated to its pro-cognitive role in normal animals and in models of ID and ASD, homozygous *PDE2A* mutations have been identified in patients affected by ID, ASD and epilepsy. To clarify this apparent paradox about the role of PDE2A in brain development, we characterized here *Pde2a^+/−^* mice (homozygote animals being not viable) at the behavioral, cellular, molecular and electrophysiological levels. *Pde2a^+/−^* females display a milder form of the disorder with reduced cognitive performance in adulthood, conversely males show severe socio-cognitive deficits throughout their life. In males, these phenotypes are associated with microglia activation, elevated glutathione levels and increased externalization of GluR1 in CA1, producing reduced mGluR-dependent LTD. Overall, our results reveal molecular targets of the PDE2A-dependent pathway underlying socio-cognitive performance. These results clarify the mechanism of action of pro-cognitive drugs based on PDE2A inactivation, which have been shown to be promising therapeutic approaches for Alzheimer Disease, schizophrenia, FXS as well as other forms of ASD.

## Introduction

Cyclic adenosine monophosphate (cAMP) and cyclic guanosine monophosphate (cGMP) are secondary messengers involved in numerous cellular pathways. They coordinate a multitude of pathophysiological processes *via* the specific activation of cAMP-dependent and cGMP-dependent protein kinases, PKA and PKG, respectively (1). The homeostasis of these secondary messengers is finely regulated both during their synthesis by adenylate cyclase and guanylate cyclase and during their degradation by members of the phosphodiesterase (PDE) superfamily of enzymes (2). Different PDEs are expressed in the brain to varying extent (1,3).

In neurons, cAMP plays a key role in synaptic transmission through short- and long-term mechanisms. cAMP acts directly on ligated-gated channel neurotransmission, neurotransmitter synthesis, storage, release, receptor sensitivity, cytoskeletal organization and remodeling, and neuronal growth and differentiation (1). cAMP is a well-known regulator of microglial function and activation (4–6), affecting microglial and macrophage polarization to pro-inflammatory or anti-inflammatory phenotypes (7). In addition, nitric oxide/cyclic guanosine monophosphate (NO/cGMP) signaling plays a key role in synaptic plasticity (8) and neuroinflammation (9). Strong evidence indicates that the cGMP/PKG pathway is involved in the modulation of glial cell activity as well as in neuroinflammatory and neurodegenerative processes (10,11).

Phosphodiesterase 2A (PDE2A) is a dual-substrate enzyme that hydrolyzes both cAMP and cGMP by its three isoforms located in the cytosol, intracellular membranes, and mitochondria, respectively. cGMP binds to the c<u>G</u>MP-binding PDE, *<u>A</u>nabaena* adenylyl cyclases, *E. coli* <u>F</u>hlAs (GAF) domain of PDE2A and induces a conformational change, resulting in enhanced hydrolysis of cAMP and cGMP (12–14). PDE2A is abundant in the brain being the most highly expressed PDE in the human hippocampus, frontal cortex, and striatum, all of which are essential for cognition and social interactions (1,15–17). PDE2A is also localized in the axons and nerve terminals of principal neurons, and is the only PDE associated with docked synaptic vesicles (18). Remarkably, about 50% of the total amount of hydrolyzed cGMP is PDE2A-dependent during rat brain synaptogenesis (19).

Inhibition of PDE2A is known to have pro-cognitive effects (20–28). Indeed, the administration of TAK-915, an inhibitor of PDE2A, to MK-801-treated rats, a schizophrenia model, reversed deficits in memory and social interaction in these animals (29). In agreement with the “cAMP theory” (30), improvement in memory and social interaction deficits were shown in Valproate-exposed rats, an environmental model of ASD, and in the *Fmr1*^-/y^ mouse and rat, rodent models of Fragile X Syndrome (FXS) (31,32) treated with another PDE2A specific inhibitor, Bay607550. Consistent with these findings, we showed increased expression and activation of PDE2A as a prominent characteristic of the pathophysiology of both these disorders (33). Homozygous mutations in *PDE2A* have been found in patients with autism spectrum disorder (ASD) and intellectual disability (ID) ranging from moderate to severe (34–37). Moreover, reduced levels of PDE2A have been reported in *post-mortem* analyses of the amygdala and the frontal cortex obtained from patients affected by schizophrenia (SCZ) (38).

To better define the role of PDE2A in neurodevelopment, we characterize here *Pde2a^+/−^* mice, since *Pde2a* knock-out in this species is lethal during embryogenesis (16). We show that these animals are a novel developmental brain disorder (DBD) model displaying socio-cognitive deficits in males during infancy, adolescence, and adulthood, while females display only a deficit in cognitive performance in adulthood. This disorder in males is also characterized by altered synaptic plasticity associated to an abnormal AMPA receptor (R) trafficking and microglia activation during pruning age.

## Results

### PDE2A levels and activity are reduced in *Pde2a^+/−^* mice

Since *Pde2a*-KO mice are not viable and die *in utero* at E15.5 (39), we decided to analyze heterozygous animals (*Pde2a^+/−^*). The levels of PDE2A in the hippocampus and cortex of both male and female mice at one month of age were lower than those in wildtype (WT) mice, as shown by Western blot (WB) analysis (Fig. 1A-B and 1C, respectively). Consistent with this result, PDE2A specific activity (measured by degradation of cGMP in the presence of Bay607550) is 35% lower in *Pde2a^+/−^*compared to WT mice at the same age (Fig. 1D). Reduced expression of PDE2A was also observed by WB in protein extracts obtained from the brains of *Pde2a^+/−^*mouse embryos at E14.5, compared to WT mice (Fig. 1E). To show the impact of reduced PDE2A levels, we measured the size of the first neurite after 2 days of neuronal culture *in vitro* (DIV). As expected from our previous studies (31), this structure is longer in *Pde2a^+/−^* neurons compared to WT neurons (Fig. 1F-G).

**Fig. 1:**
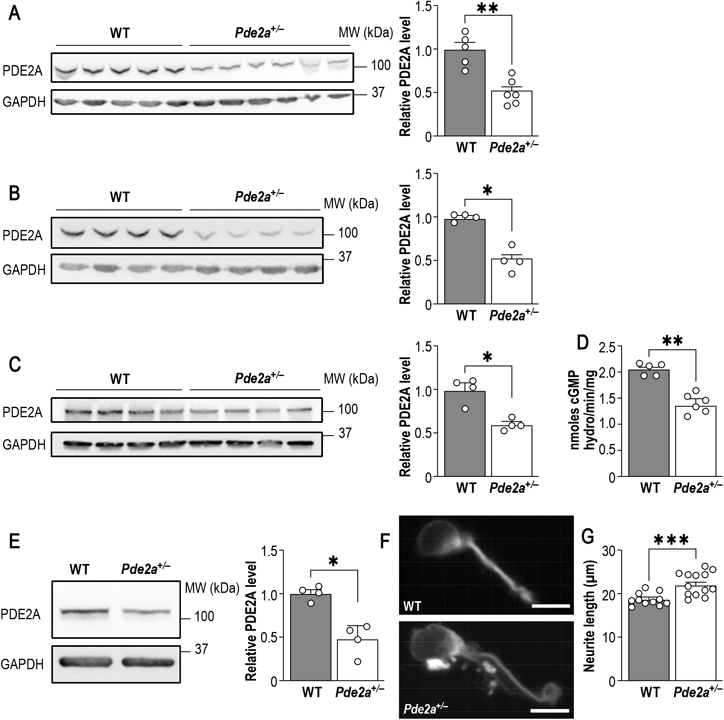
PDE2A levels are correlated with neurite growth regulation. **(A,B)** Representative Western Blot and quantification show reduced PDE2A levels in one-month male *Pde2a^+/−^* hippocampus and cortex (n=4-6). **(C)** Representative Western Blot and quantification show reduced PDE2A levels in one-month female *Pde2a^+/−^* cortex (n=4). **(D)** Specific PDE2A activity, measured by degradation of cGMP, is significantly reduced in *Pde2a^+/−^* compared to WT mice (n=5-6). **(E)** Representative Western Blot and quantification show reduced PDE2A levels in *Pde2a^+/−^* cortical neurons at E14.5 compared to WT neurons (n=3-4). **(F)** Representative pictures of 2 days *in vitro* cultured WT and *Pde2a^+/−^* primary cortical neurons (scale bar: 10 µm). **(G)** Histogram of neurite length showing longer neurite of *Pde2a^+/−^*neurons (n=11-13). The data are represented as means ± SEM and analyzed using a Mann-Whitney test (adjusted p value: *P<0.05, **P<0.01, ***P<0.001).

### *Pde2a^+/−^* mice display deficits of socio-cognitive behaviors at infancy, adolescence and adulthood

We first performed a behavioral characterization of *Pde2a^+/−^*male mice. Ultrasonic vocalizations (USVs) emitted by rodent pups in response to separation from the mother and nest play an essential communicative role in mother-offspring interactions (40). Isolation-induced USVs were collected for 3 min as social communication signals from mouse pups on postnatal day (PND) 10. *Pde2a^+/−^* pups emitted significantly fewer USVs than WT *Pde2a^+/+^* littermates (Fig. 2A). The homing behavior test measures the tendency of rodent pups to maintain body contact with dams and siblings by discriminating the nest odor from a neutral odor (40). At PND 13, no differences between *Pde2a^+/−^* and WT were found (Fig. 2B). Body weight was measured to ensure that the reduced USVs were not the result of being physically smaller, because body weight is known to alter pup USV emissions (41). However, weight did not differ between the genotypes at 10 and 13 PND (Supp. Fig. 1A-B), indicating normal growth in *Pde2a^+/−^* mice. Adolescent male *Pde2a^+/−^*mice showed a reduced frequency of social interactions compared with their WT littermates (Fig. 2C). Overall, it appears that physiological PDE2A activity is critical for specific social behaviors, as *Pde2a^+/−^* pups displayed early communicative deficits but normal social discrimination ability and adolescent animals showed reduced social interactions. Anxiety-like behavior was evaluated using dark-light (DL) and elevated zero maze (EZM) tasks. During the DL test, *Pde2a^+/−^* adult mice did not display behaviors different from those of their WT littermates (Supp. Fig. 1C). In the EZM task, *Pde2a^+/−^* mice displayed a higher latency to exit from the closed arm of the maze (Fig. 2D, left panel), although the time spent in the open arms did not differ from that of the WT controls (Fig. 2D, right panel). These behaviors do not suggest the presence of anxiety-like behaviors in this model but rather a delay in adaptation to novelty.

**Fig. 2:**
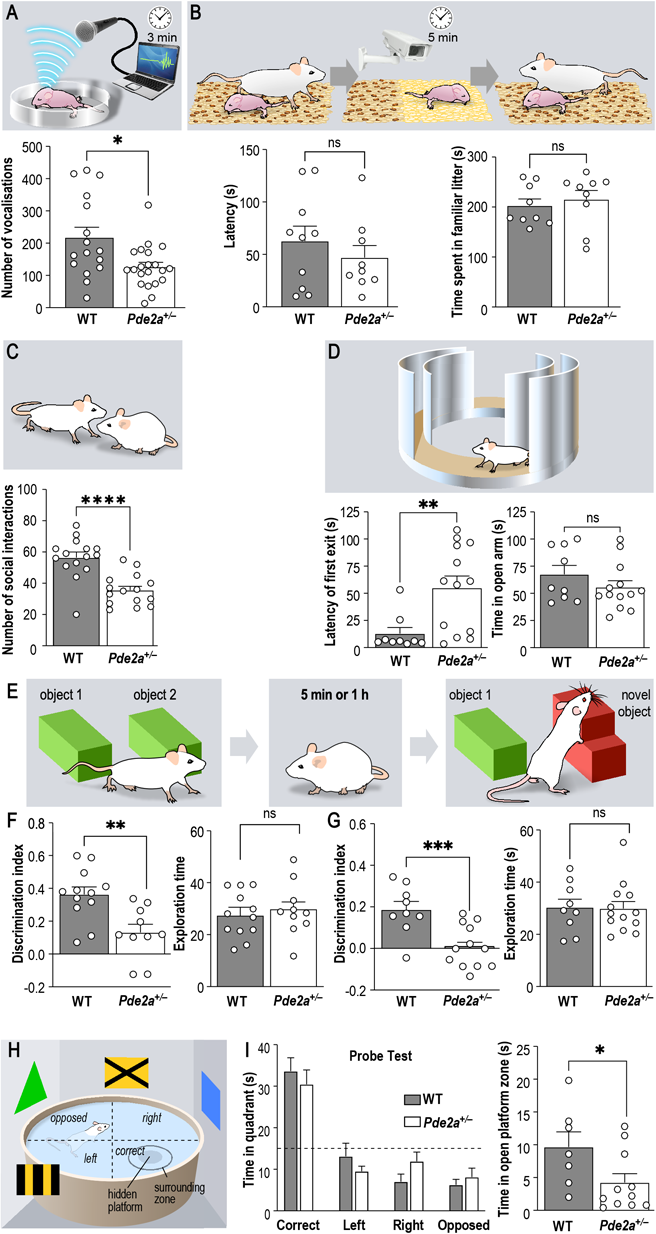
*Pde2a^+/−^* mice displayed deficits in various socio-cognitive tasks at different ages. **(A)** *Pde2a^+/−^* pups emit significantly less ultra-sonic vocalisations (USVs) when removed from the nest at P10 (n=16-21). **(B)** *Pde2a^+/−^*pups show no deficits in the homing test compared to WT pups at P13, as latency to reach the familiar litter and the time spent in it are not different (n=9-10). **(C)** At one-month age, adolescent *Pde2a^+/−^* mice make significantly less interaction when presented to a new mouse compared to WT (n=15). **(D)** At two-month age, adult *Pde2a^+/−^* mice are significantly slower to first exit the closed arm from the zero-maze apparatus, but spent the same amount of time in open arms at the end compared to WT mice (n=9-13). **(E)** *Pde2a^+/−^* mice show significantly lower preference for a novel object than WT mice, with a diminution of discrimination index, whether with 5 minutes **(F)** or 1 hour **(G)** between learning and test phase (n=10-12). **(H,I)** *Pde2a^+/−^* mice showed a slight deficit in spatial memory, as they spent significant less time in open platform zone **(I right)** during the Probe test of Morris Water Maze (n=7-11). The data are represented as means ± SEM and analysed using Mann-Whitney test and **(I left)** Two-Way ANOVA (p value: *P<0.05, **P<0.01, ***P<0.0005, ****P<0.0001).

To assess learning and memory in those animals, we first investigated their working and short-term memories using a novel object recognition (NOR) task. We performed the test in two different sessions, adjusting the time between the learning and test phases to 5 min and 1 h, respectively (Fig. 2E). In both settings, *Pde2a^+/−^* mice showed a significantly reduced discrimination index compared to WT littermates (Fig. 2F-G, left panel), while the exploration time was the same for WT and *Pde2a^+/−^* mice during training (Supp. Fig. 1D-E), and during the test phase (Fig. 2F-G, right panel). We assessed spatial memory in *Pde2a^+/−^*mice by using the Morris Water Maze task (Fig. 2H). In the Cue phase (learning of the hidden platform’s location with the help of visual cues), *Pde2a^+/−^* mice reached the visible platform at a significantly slower pace than WT mice on the first day of the experiment. During the second day, they had the same performance as WT mice (Supp. Fig. 1F, left panel). During both the learning and probe phases, the performances of *Pde2a^+/−^* mice were comparable (Supp. Fig. 1F, right panel; Fig. 2I, left panel). However, during the probe phase, *Pde2a^+/−^* mice spent significantly less time in the exact location of the platform (Fig. 2I, right panel) and in its surrounding zone. Collectively, the results of these tests showed that *Pde2a^+/−^* male mice display deficits in working and short-term memory and, to a lesser extent, in spatial memory. Moreover, the behavior of these mice in the cue phase reinforces our interpretation that they have a delay in adaptation to novelty.

We studied animal activity using actimetry and observed that mice of both genotypes crossed the beams detecting movement with the same frequency during the 68 h of the task (Supp. Fig. 1G). During the same time, we observed that the number of rearings was not significantly different between WT and *Pde2a^+/−^* mice (Supp. Fig. 1H), and animals of both genotypes had the same performance in the rotarod task (Supp. Fig. 1I).

Afterwards, we analyzed the behavior of *Pde2a^+/−^* female mice. We did not find any abnormalities in the isolation-induced USV emissions or discrimination abilities in the homing test (Supp. Fig. 2A-B). We measured a small but significant decrease of social interactions in adolescent *Pde2a^+/−^*females (Supp. Fig. 2C), which was normalized to the WT levels at the adult age (Supp. Fig. 2D). No differences between the genotypes were found in the EZM task (Supp. Fig. 2E). In the NOR task, *Pde2a^+/−^* females showed a significantly lower discrimination index than their WT littermates (Supp. Fig. 2F), whereas the exploration time during the learning and test phases was unaffected (Supp. Fig. 2G-H). In conclusion, *Pde2a^+/−^*females display only mild memory deficits in adulthood. No spontaneous epileptic crisis was observed in *Pde2a^+/−^* mice of both sexes.

Considering the mild phenotype in females, we decided to focus only on males to understand the molecular and cellular consequences of the reduction in PDE2A.

### PDE2A genetic reduction results in multiple cellular abnormalities

We employed Golgi staining to study the morphology of dendritic spines in the brain of male *Pde2a*^+/−^ mice. In the hippocampus, the length of dendritic spines displayed a small but statistically significant increase compared to that in the WT, while the number of spines did not differ between the two genotypes (Fig. 3A). Conversely, the length and number of dendritic spines in the cortex were reduced compared to those in the WT (Fig. 3B). This could be explained by an altered pruning process, which peaks two weeks after birth and is strongly correlated with microglial activity (42,43). cAMP is a well-known regulator of microglial function and activation (4–6) and nitric oxide/cyclic guanosine monophosphate (NO/cGMP) signaling plays a key role in microglia activity and neuroinflammation (9–11). In two-week-old *Pde2a^+/−^* mice, the number of cells expressing the microglia marker IBA1 was increased in the cortex compared to WT animals; however, it was normal in the hippocampus (Fig. 3C, Supp. Fig. 3A). We confirmed these findings by Western blot, which showed an increased level of IBA1 in the cortex of two-week-old *Pde2a^+/−^* mice compared to that in WT mice (Supp. Fig. 3B), but not at one month (Supp. Fig. 3C). For further characterization, we studied the morphology of microglia in the cortex (Fig. 3D). Microglia are highly plastic cells whose morphological modifications indicate deregulation of their homeostasis. Confocal imaging revealed that the total length and number of filaments were increased in *Pde2a^+/−^* cortex compared to those in the WT (Fig. 3E-F). In addition, these cells were more ramified than the WT microglia (Fig. 3G), as assessed by Sholl analysis(44). Using this approach, we also found that the number of microglia cells was higher in the cortex of *Pde2a^+/−^* mice (Supp. Fig. 3D), which confirmed our observations (Fig. 3C).

**Fig. 3:**
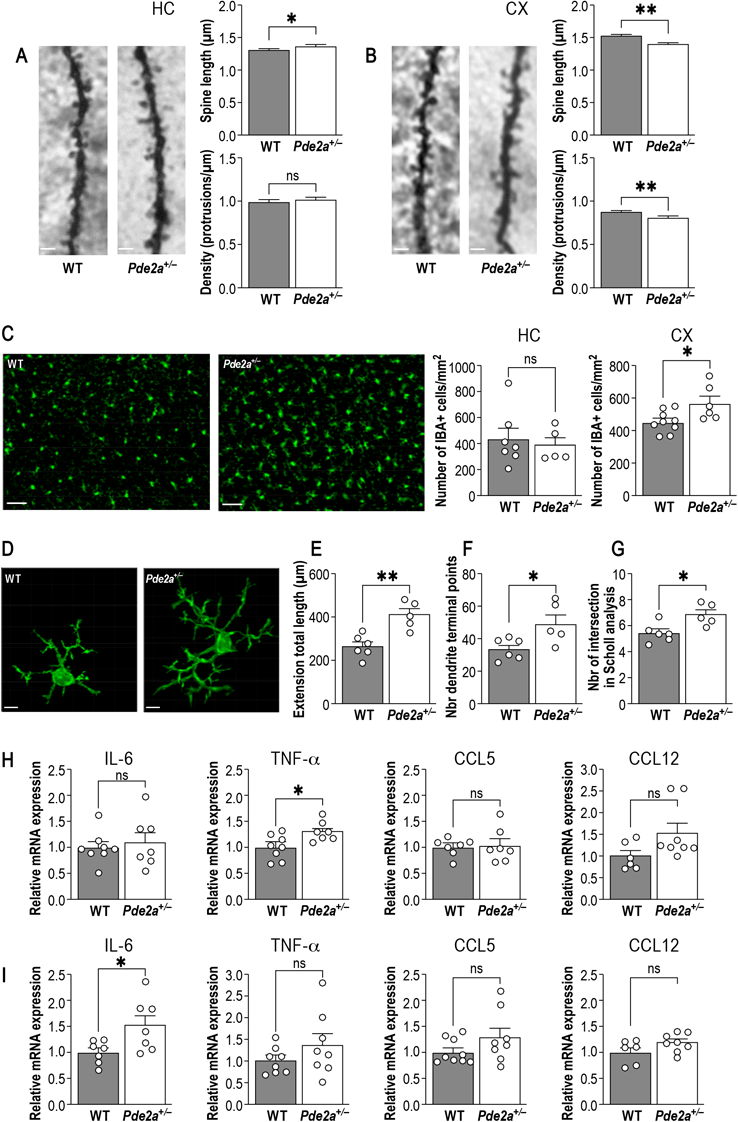
*Pde2a^+/−^* mice present dendritic spine and microglia morphology alteration, associated with pro-inflammation markers. (A) Dendritic spines are significantly longer in CA1 of *Pde2a^+/−^* hippocampus while spine density is not different compared to WT. **(B)** Dendritic spine length and density are significantly reduced in the cortex of one-month *Pde2a^+/−^* mice (bar scale: 2µm). **(C)** Representative image of Iba1 positive cells in WT and *Pde2a^+/−^* cortex at P13 (bar scale : 25µm) showing that number of Iba1 positive cells in hippocampus are not different between WT and Pde2a+/− mice (n=5-7), while in cortex *Pde2a^+/−^* mice have significantly more IBA1+ cells (n=6-9). **(D)** Representative pictures of IBA1 immunoreactive microglia in WT and *Pde2a^+/−^*cortex at P13 (bar scale : 4µm). (E) *Pde2a^+/−^* microglia present significantly higher extension total length, **(F)** more dendrite terminal points **(G)** and is wider, as it crossed more Scholl intersections (n=5-6). **(H,I)** Relative mRNA expression of various inflammatory markers in hippocampus and cortex of WT and *Pde2a^+/−^*mice at P13 (n=6-9). The data are represented as means ± SEM and analyzed using Mann-Whitney test. (p value : *P<0.05, **P<0.01)

Since microglial activation is one of the earliest features of nearly any change in brain physiology (45), we further investigated the abundance of cytokines and chemokines in the cortex and hippocampus using qPCR. TNF-α and IL-6, two pro-inflammatory cytokines, were upregulated in two-week-old *Pde2a^+/−^*mice compared to WT mice, in the hippocampus and cortex, respectively (Fig. 3H-I), whereas CCL5 and CCL12 were not differently expressed in both regions. In four-week-old *Pde2a^+/−^* mice, TNF-α and IL-6 expression was increased in the hippocampus (Supp. Fig. 3E), as opposed to what was observed in the cortex, where only CCL5 expression was significantly higher (Supp. Fig. 3F). However, CCL5 was not differentially expressed in the hippocampus, and CCL12 was not expressed in the cortex at this age (Supp. Fig. 3E-F).

### Mitochondria and metabolism

A specific PDE2A isoform is localized in mitochondria (13) where it plays an important role in defining the morphology of those organelles (13,46). Consistent with these findings, the presence of numerous thick and irregular mitochondria was reported in the fibroblasts of patients carrying a mutation in the *PDE2A* gene (34). We studied the morphology of mitochondria using Transmission Electron Microscopy (TEM) images in one-month old WT and *Pde2a^+/−^* hippocampus and cortex (Fig. 4A). We observed that *Pde2a^+/−^* mitochondria were more electron-dense and their area was significantly decreased compared to WT in both the cortex and hippocampus (Fig. 4B-C).

**Fig. 4:**
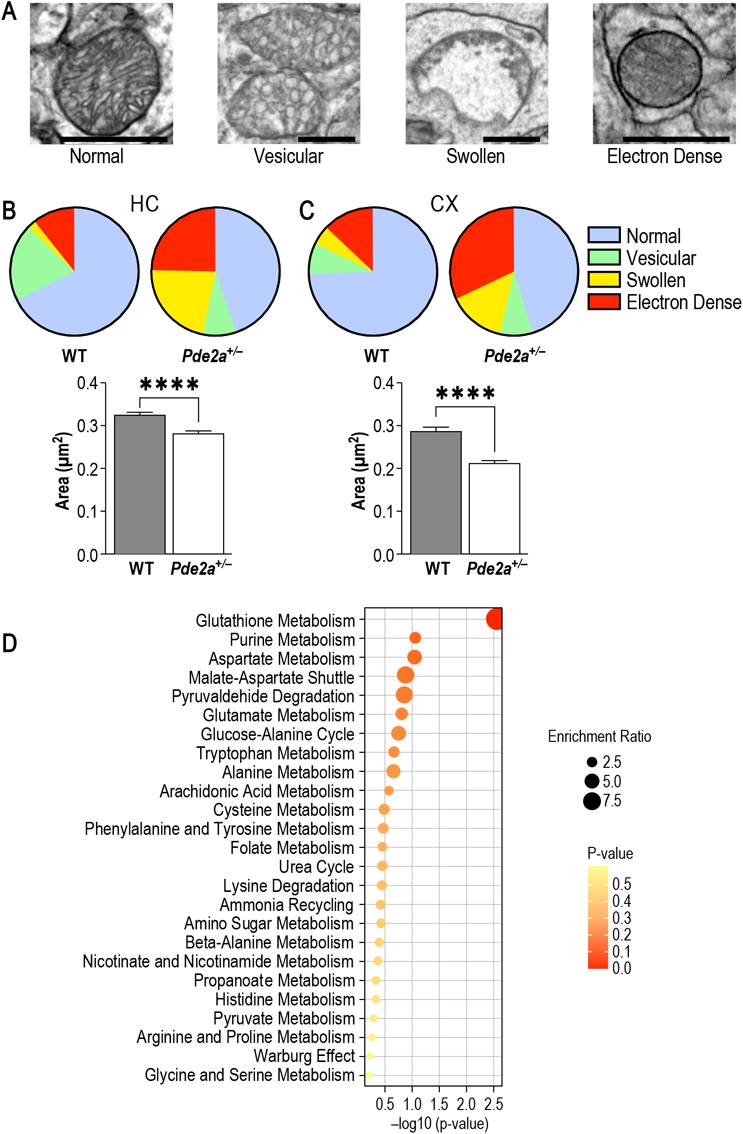
*Pde2a^+/−^* mice have abnormal mitochondria associated with metabolomics changes. **(A)** Representative TEM images of four main mitochondrial morphology types (bar scale: 1µm) **(B,C)** One-month-old *Pde2a^+/−^* mice presents abnormal mitochondria proportion and their mitochondria are significantly smaller in both hippocampus and cortex compared to WT mice. **(D)** Enrichment overview of metabolism pathways dysregulated in cortex of *Pde2a^+/−^* mice. The size and color of the dots indicate the enrichment ratio and the level of P value, respectively.The data are represented as means ± SEM and analyzed using the Mann-Whitney test (p value: ****P<0.001)

Considering the critical role of mitochondria in cellular metabolism (47), and the strong association existing between mitochondrial dysfunction and neuroinflammation (48), we performed untargeted metabolomics of cortex extracts from WT and *Pde2a^+/−^* mice. The levels of L-tryptophan, a key amino acid implicated in serotonin synthesis, glutamate and of the related compounds gamma-glutamyl-glutamic acid, L-pyroglutamic acid, N-acetyl-aspartylglutamic acid and glutathione were increased in the cortex of *Pde2a^+/−^* compared to WT mice (Supp. Table 1). Glutathione metabolism was the most dysregulated pathway in the *Pde2a^+/−^*mice cortex (Fig. 4D and Supp. Table 2).

### AMPA-dependent mGluR-dependent Long-Term Depression is disrupted in *Pde2a^+/−^*mice

We previously showed that the pharmacological inhibition of PDE2A by Bay607550 reduces the exaggerated mGluR-dependent Long-Term Depression (LTD) in the *Fmr1*-KO mouse CA1 (31). At that time, we did not investigate the mechanism of action of PDE2A in the modulation of this synaptic plasticity, which can be now explored by using the *Pde2a^+/−^* mice. First, the hippocampal mGluR-dependent LTD was investigated in CA1 of WT and *Pde2a^+/−^* mice. Extracellular AMPA receptor-mediated field excitatory postsynaptic potentials (fEPSPs) were evoked from CA1 region of hippocampus after electrical stimulation of CA1. Bath application of DHPG, an agonist of group I mGluRs, (100 µM, 5 min) induced a mGluR-LTD that is weaker in *Pde2a^+/−^* mice when compared with WT mice (Fig. 5A-B). Since DHPG action induces the internalization of GluR1 *via* DARPP-32-PP1-dependent dephosphorylation of Ser845 (49), we measured the ratio of surface/total GluR1 in hippocampal slices of four-week-old mice and we found that GluR1 receptors are localized at the membrane surface at higher levels in *Pde2a^+/−^* slices compared with WT after DHPG stimulation (Fig. 5C-D). This suggests an elevated abundance of GluR1 at the membrane upon DHPG. On the other side, it is known that the phosphorylation of is Ser845-GluR1 is cAMP/PKA dependent. We tested the level of GluR1 phosphorylation through WB using an antibody to monitor the phosphorylation of Ser845-GluR1 in hippocampus extracts of four-week-old WT and *Pde2a^+/−^* mice. We found increased phosphorylation of GluR1 (Fig. 5E-F) indicating an increased membrane abundance of GluR1 also at the steady-state condition in *Pde2a^+/−^* mouse hippocampus and, consequently, an altered responsiveness of AMPA receptor (R).

**Fig. 5:**
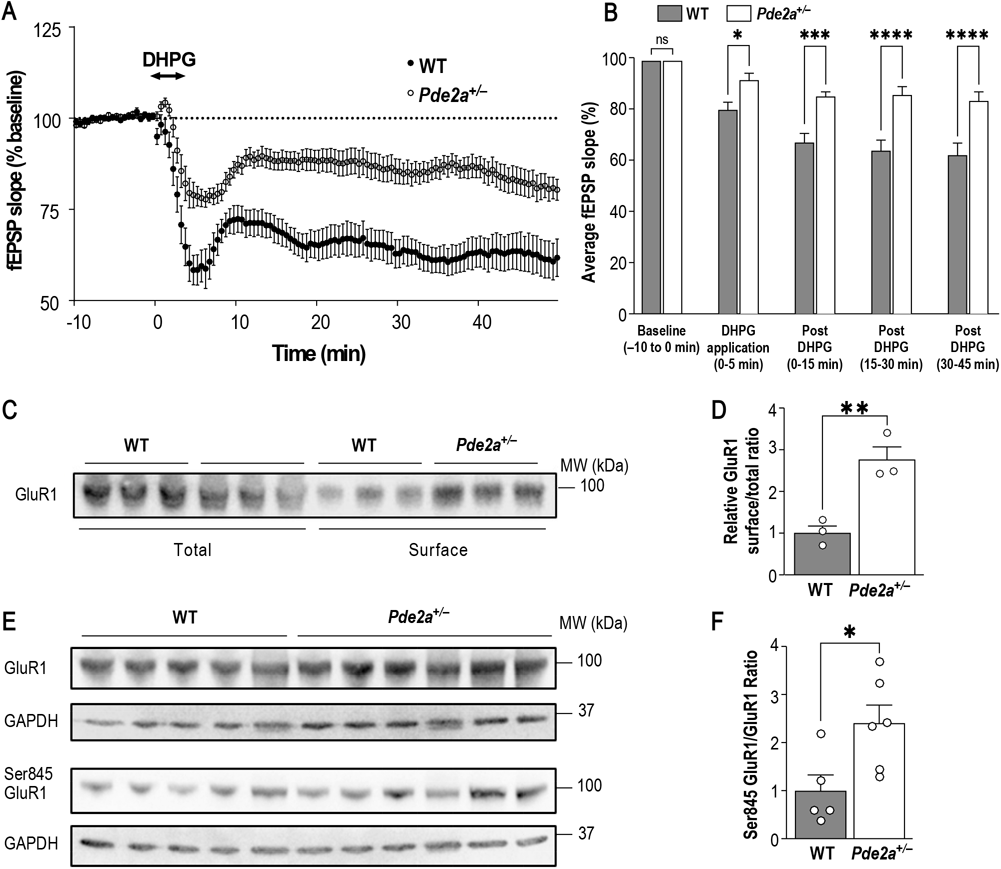
Hippocampal mGluR-LTD is impaired in *Pde2a^+/−^* mice. **(A)** DHPG (100 μM, 5 min) induced a mGluR-LTD of fEPSPs recorded from CA1 pyramidal neurons obtained from WT and *Pde2a^+/−^* mouse slices (n = 10). CA1 LTD induction was weaker in *Pde2a^+/−^* mice when compared to WT mice. **(B)** Bar graphs represent the average slope response during baseline (−10 to 0 min), DHPG application (0-5 min), 0-15 min after DHPG application, 15-30 min after DHPG application, 30-45 min after DHPG application. **(C)** Western Blot and **(D)** quantification show increased GluR1 presence in surface after DHPG application in one-month *Pde2a^+/−^*hippocampus (n=3). **(E)** Western Blot and **(F)** quantification show an increase phosphorylation ratio of GluR1 in *Pde2a^+/−^* hippocampus compared to WT (n=5-6). GluR1 and Ser845 GluR1 were previously normalized by their respective GAPDH level. The data are represented as means ± SEM and analysed using the Mann-Whitney test and t test for **D** (p value: *P<0.05, **P<0.01, ***P<0.0005).

In summary, our results are consistent with an increased PDE2A-dependent abundance of cAMP impacting AMPAR trafficking.

## Discussion

### A sex-dependent phenotype

Mutations in *PDE2A* have been associated with DBD such as ID, ASD, epilepsy, and movement disorders in homozygous conditions (37). Only a few cases have been described; in some patients, the first neuropsychiatric evaluation was performed during adolescence or later (34–37). We present here the first characterization of *Pde2a^+/−^* mice as a model of this disorder since homozygote *Pde2a* mutant mice are embryonically lethal due to liver (50) and heart defects (39). *Pde2a^+/−^*mice displayed socio-cognitive deficits, as we assessed using multiple behavioral tasks. In particular, male *Pde2a^+/−^* pups displayed early communicative deficits at P10 and reduced social interaction at P30. Female *Pde2a^+/−^* mice did not display these phenotypes at infancy and showed mild social deficits during adolescence, that were rescued in adulthood, indicating that the phenotype was mild and transitory. It is important to note that animals of both sexes exhibited memory impairments at P60 in the novel object recognition task. These data confirm the role of PDE2A in cognition, as previously suggested (1), but strongly highlight the role of this protein in male social interaction. It is then possible to hypothesize that there is a sex-dependent regulation of PDE2A, as is the case with another PDE, PDE4D, which is differentially modulated in the male brain compared to the female brain (51). *Pde2a^+/−^* female mice display a severe phenotype only at adulthood, which might result from sex-specific compensation of *Pde2a* partial inactivation. Similarly, the partial loss of *Gabrb3* differentially alters cerebellar physiology in a sex-specific manner, leading to a sex-dependent compensatory mechanism in young male mice that is not maintained in adults (52). Considering our present data, we can conclude that partial inactivation of *Pde2a* in mice generates two different disorders in males and females.

No epileptic crises or abnormal movement disorders were observed in infant, adolescent, or young adult *Pde2a^+/−^* animals of both sexes. This is different from what was described in patients carrying homozygote mutated forms of *PDE2A*. A rotarod test performed at 6 months did not show any phenotype in male or female *Pde2a^+/−^* mice. The fact that homozygous mutations are not lethal in patients suggests the presence of residual activity of PDE2A in these individuals, or, more likely, that its total absence is compensated for in the human liver and heart by the activity of another PDE, but this is not the case in mice. This compensation could also be present in the brains of those patients, since heterozygote individuals are not described as being affected by ASD or ID, indeed no neuropsychiatric evaluation was reported for the patients’ parents carrying a heterozygous *PDE2A* mutation. However, only limited functional analyses of PDE2A mutant proteins identified in patients have been performed, and in some cases a poor description of the genetic variants has been reported. The endogenous activity of PDE2A was not measured in these patients (34–37). Consequently, we could not assess whether the variants identified thus far were gain- or loss-of-function mutations. Interestingly, even if we can consider the characterization of patients carrying mutations in *PDE2A* as preliminary, we noticed that for two female patients a regression of the neurodevelopmental phenotype was described (35), leading to the possible speculation that in these patients a sex-specific phenotype is detectable in early infancy.

### *Pde2a^+/−^* mice: cellular and molecular phenotypes in brain

In the analysis of *Pde2a^+/−^* mice, we focused on the role of PDE2A in hippocampus and cortex, the two brain regions where the translation of its mRNA is negatively modulated by FMRP (1,31,33). This reason, together with the mild neurodevelopmental phenotype of female *Pde2a^+/−^* mice, pushed us to carry out the molecular and cellular characterization of the brain lacking PDE2A only in male animals.

### Microglia

A large body of literature has reported the implications of cAMP and cGMP in neuroinflammation (4–6). We found an increased number of IBA1+ cells associated with altered cell morphology only in the cortex from two-week-old *Pde2a^+/−^* mice, but no differences were observed in the hippocampus of the same mice. Increased expression levels of IL-6 and TNF-α were present in both the cortex and hippocampus of *Pde2a^+/−^* mice during development. Elevated expression of these cytokines is associated with ASD and schizophrenia (53,54). Furthermore, the altered morphology of microglia observed in the cortex during the developmental pruning peak may explain the reduced number of dendritic spines observed in this brain region in one-month-old mice. Surprisingly, at one month of age, when we did not observe altered IBA1 expression, the expression level of IL-6 and TNF-α of *Pde2a^+/−^* mice was only increased in hippocampus but not in the cortex, where, conversely, we observed an increased abundance of CCL5. Interestingly, this chemokine has been described as a neuromodulator and its elevated expression has been found in patients with ASD (70, 71,72). This result strongly suggests the need of to study the expression level of cytokines and chemokines during developmental age to monitor microglia plasticity in DBDs.

### Mitochondria and metabolism

Mitochondria are indispensable organelles that play a crucial role in cellular energetics, metabolism, and survival (47). Accumulating evidence indicates that aberrant mitophagy, respiratory chain deficits, and oxidative stress resulting from mitochondrial dysfunction lead to aberrant neurodevelopment (57). Neurological disorders associated with mitochondrial dysfunction are often associated with epilepsy in children. Oxidative damage induces hyperexcitability by modifying the excitation/inhibition balance (58). This may contribute to epileptic conditions in patients carrying homozygous mutations in *PDE2A*.

It has been reported that the mitochondrial isoform of PDE2A (PDE2A2) is mostly localized in the inner mitochondrial membrane and regulates mitochondrial fusion/fission (13) and mitophagy (46). All these findings seem consistent with our data showing that the total mitochondrial area in the *Pde2a^+/−^*hippocampus and cortex is smaller and an increased number of abnormal mitochondria is observed in these tissues.

Mitochondrial dysfunction is also strongly associated with neuroinflammation (59). Microglial immune functions have a high energy demand, which is regulated by mitochondria (60). Therefore, mitochondrial quality is important for cellular homeostasis (61). Interestingly, the most deregulated metabolites in the cortex of *Pde2a^+/−^* mice (glutathione and purine metabolism) are well known to contribute to oxidative stress (62). Recent studies have suggested that mitochondrial dysfunction, including mitochondrial DNA (mtDNA) damage, metabolic defects, and quality control disorders, precedes microglial activation and subsequent neuroinflammation (63). In this context, mitochondrial alterations may be associated with microglial activation.

### GluR1 trafficking and mGluR-dependent LTD

We have shown an increased phosphorylation level of S845-GluR1 in *Pde2a*^+/−^ hippocampus. Even if S845-GluR1 phosphorylation is known to be PKA-dependent (49,64,65), the implication of a specific PDE was to our knowledge never shown before in hippocampus. Overall, our results clearly demonstrate the critical role of PDE2A in hippocampal AMPAR trafficking and suggest an abnormal responsiveness of AMPAR in the hippocampus of these animals. Indeed, upon DHPG stimulation, *Pde2a*^+/−^ CA1 neurons display an increased externalization of GluR1 and a reduced mGluR-dependent LTD. It has also been shown that mGluR5 activation promotes GluR1 Ser- 845 phosphorylation via inactivation of protein phosphatase-1 (PP1) (66), which targets GluR1 Ser-845. Thus, our results suggest that in the presence of an elevated level of GluR1 Ser-845 phosphorylation (50% increase compared to WT-Fig. 5E-F) the inhibition of PP1 maintains elevated Ser-845 phosphorylation compared to WT with a consequent reduced AMPAR internalization and an impaired mGluR-dependent-LTD.

### Conclusions

*Pde2a^+/−^* mice represent a new DBD model and a useful tool for understanding the altered trajectory of brain development in pathological conditions. We highlight here that PDE2A-altered levels have a negative impact on the socio-cognitive phenotype, both when increased and reduced, strongly suggesting that PDE2A is a Goldilocks factor. Multiple molecular and cellular phenotypes characterize the brain of these mice. Remarkably, similar to *Pde2a^+/−^* mice, other forms of DBDs, such as Phelan-McDermid (SHANK3) (67,68), Tuberous Sclerosis (TSC1/TSC2) (69,70), Cowden (PTEN) (71), mTORC2 (72), display an impaired mGluR5-dependent LTD. Consistent with our data, inhibition of *Tsc2*, *PTEN* or *mTORC2* resulted in increased PKA activity and the presence of longer neurites (73,74). This finding suggests an involvement of PDE2A in the pathophysiology of these disorders.

Our results, contributing to deciphering the mechanism of action of PDE2A, also allow to make hypotheses about the specific mechanism of action of pro-cognitive drugs based on PDE2A inhibition (1,75), that have been proposed in preclinical studies to treat Alzheimer Disease (AD), Schizophrenia, ASD and FXS (1,19).

## Methods

### Animals

The experiments were performed following the Animals in Research: Reporting *In Vivo* Experiments (ARRIVE) guidelines and the European Community Directive 2010/63/EU. The experiments were approved by the local ethics committee (Comité d’Ethique en Expérimentation Animale CIEPAL-AZUR N. 00 788.01; APAFIS #39606-202110061015176 v8) and by the French Ministry of Research. WT and *Pde2a^+/−^* mice were on a C57Bl/6J background. Mutant mice (B6; 129P2-Pde2a <tm1Dgen>/H EM: 02366) were obtained from EMMA, UK (16) and were revitalized at Plaisant SRL (Rome, Italy). All animals used in behavioral experiments were littermate and were generated and housed in groups of six in standard laboratory conditions (22°c, 55 ± 10% humidity, 12-h light/12-h dark diurnal cycles), with food and water provided *ad libitum*. Mice were weaned at the third postnatal week, chipped and genotyped by PCR at the end of the experiments. *Pde2a^+/−^* mice and WT alleles were detected by PCR assay in which three primers (see Table S3) amplify a 247 bp fragment in WT and two fragments in *Pde2a^+/−^*, the same one as in the WT and an additional 354 bp fragment.

### Neuronal Cultures and Neurite Length Measurements

Primary cortical neurons were prepared from embryos at E14.5 obtained from pregnant C57Bl/6J *Pde2a^+/−^* mice, as previously described (31). After 48h and fixation, neurons were labeled with anti-Tuj1 antibody and the size of individual growing neurites was manually measured using the Fiji software (https://fiji.sc), from the soma to the end of the neurite.

### PDE2A activity assay

Half brains of 1-month old mice were collected and snap frozen in liquid nitrogen and stored at −80°C. Frozen brains were then homogenized using a glass homogenizer (15 strokes, 4°C) in 20 mM Tris-HCl buffer pH 7.2 containing 0.2 mM EGTA, 5 mM β-mercaptoethanol, 2% v/v antiprotease cocktail (Sigma–Aldrich), 1 mM PMSF, 5 mM MgCl_2_, 0.1% v/v Triton X-100. The homogenates were centrifuged at 14.000*g* for 30 min at 4°C.

PDE activity was measured in the supernatant with the method described by Thompson and Appleman (76) in 60 mM Hepes pH 7.2, 0.1 mM EGTA, 5 mM MgCl_2_, 0.5 mg/ml bovine serum albumin (BSA), and 30 μg/ml soybean trypsin inhibitor, in a final volume of 0.15 ml. To evaluate the enzymatic activity of PDE2A, the specific inhibitor Bay 607550 was added to the reaction mix at a final concentration of 0.1 μM, before the addition of the substrate. PDE2A activity was calculated as the difference between the total hydrolytic activity and the residual hydrolytic activity assayed in the presence of the specific inhibitor (32)

### Behavioral Tasks description

**See Supplementary Information**

### Electrophysiology

**See Supplementary Information**

### Protein Extraction and Western Blot

Protein extraction was carried our as previously described (33). Samples were boiled at 95°C, or not boiled when testing for GluR1 expression. Samples were separated on NuPAGE Bis-Tris 4%–12% gels in MOPS buffer. Separated proteins were transferred to nitrocellulose membranes (Bio-Rad). Membranes were blocked with PBS-Tween (0.1%) and milk (5%), and incubated with primary antibodies overnight. Anti-GAPDH (Millipore, 1:5000), anti-PDE2A (Abcam, 1:1000), anti-GluR1 (Cell Signaling, 1:1000) and anti-Ser845-GluR1 (Cell Signaling, 1:1000) were used here. After secondary antibody incubation, membranes were revealed using Immobilon Western (Millipore). Protein bands intensity were quantified using ImageJ.

### Biochemical Measurements of Surface GluR1

Hippocampal slices (300 μm) were prepared from one-month mice and immediately placed in ice-cold ACSF saturated with 95% oxygen (O_2_) and 5% carbon dioxide (CO_2_). Slices (8 to 10 per group) were equilibrated at room temperature for 2 hours and then incubated with DHPG ((S)-3,5-Dihydroxyphenylglycine) (100 μmol/L) for 5 minutes. All treatments were carried out in the presence of D-AP5 (50 μmol/L). Drugs were removed by three washes in ACSF containing D-AP5 and left in the same buffer for an additional 40 minutes. Slices were washed once with ice-cold aCSF (5 min) and then incubated with Sulfo-NHS-SS-biotin (Pierce Chemical Company, Rockford, Illinois; 1 mg/mL in aCSF) for 30 minutes on ice. Excess of biotin was removed by two brief washes with 100 mmol/L glycine (in aCSF for 5 min) and two ACSF washes. Slices were then homogenized in the same lysis buffer, sonicated, and centrifuged at 14000*g* for 15 minutes to remove nuclear material and cell debris. Protein concentration was determined by using the Bradford assay (Bio-Rad). Sixty micrograms of proteins were used to measure total GluR1 and sixty micrograms of biotinylated proteins were incubated with streptavidin beads (Pierce Chemical Company) on a head-over-head shaker overnight at 4°C. Beads were washed three times with lysis buffer; bound proteins were eluted with loading buffer by boiling for 5 minutes. Total proteins and isolated biotinylated proteins were analyzed by Western blotting.

### Transmission Electronic Microscopy

Mice were intracardially perfused with 10 ml of 2.5% glutaraldehyde in 0.1 M cacodylate buffer. Fixed mice brains were sliced (200 µm) on a vibratome. Small sections from the cortical regions and CA1 hippocampus were microdissected under binoculars and post-fixed 2 hours in reduced 1% osmium tetroxide with 1% potassium ferrocyanide in 0.1 M cacodylate buffer to enhance the staining of membranes. Samples were rinsed in distilled water, dehydrated in increasing concentrations of acetone and lastly embedded in epoxy resin (EPON). Contrasted ultrathin sections (70 nm) with uranyl acetate and lead citrate were analyzed under a JEOL 1400 transmission electron microscope operating at 100 kV and mounted with a Morada Olympus CCD camera.

### RNA extraction and RT-qPCR

Total RNA was extracted from 20 mg of frozen cortex and hippocampus using the RNeasy mini kit (Qiagen, #74104) according to the manufacturer’s instructions. RNA was resuspended in 30 μL of nuclease-free H_2_O and treated with Turbo DNAse (Thermofisher Scientific). For each experimental sample, 1 μg of RNA was added to the RT reaction, that was performed using the SuperScript IV synthesis kit (Invitrogen). Initial amplification was performed with a denaturation step for 5 min at 65°C, followed by oligo(dT) annealing for 10 min at 23°C, primer annealing for 10 min at 53°C, and primer extension for 10 min at 80°C. Upon completion of the cycling steps, the reactions were stored at −20°C. Quantitative PCR (RT-qPCR) was performed on a light cycler 480 (Roche) with MasterMix SYBR Green (Roche) following the manufacturer’s instructions. Primer sequences are listed in Table S3.

### Immunohistochemistry experiments

Mice were perfused intra-aortically with 4% PFA and brains were collected. Immunostaining was performed on 40-μm-thick brain floating sections of 13 days old pups. Rabbit anti-mouse IBA1 (1:300, #CP290A, Biocare Medical) and goat anti-rabbit secondary antibody conjugated with Alexa 594 (1:500, Molecular Probes, USA) were used. Sections were mounted in Vectashield solution (H-1000, Vector Laboratories). Images mosaics for cell counting IBA1 positive cells were acquired with an inverted microscope for epifluorescence (Axiovert 200 M, Zeiss, Germany) through a 10X/0.45 objective. High-resolution images for morphometric analysis for IBA1 immunostaining were acquired with a Laser Scanner Confocal Microscope (TCS, SP5, Leica, Germany) through a 63X/1.4oil immersion objective with a z-step of 0.5 µm and a pixel size of 70 nm.

### Image analysis

For analysis of mitochondrial morphology, ImageJ was used to measure manually the area of each clearly visible and distinguishable mitochondrion from three different mice per genotype. For measurement of microglial morphology, we used high resolution *z*-stack of cortex images using the Imaris software (Bitplane). After a surface rendering of each channel, we analyzed the morphology using the “Filament” plug-in.

### Statistical Analysis

Results are expressed as mean ± standard error of the mean (SEM). All statistical analyses were based on biological replicates. Appropriate statistical tests used for each experiment are described in the corresponding figure legends. All statistical analyses were carried out using GraphPad Prism v.10.0.

## Acknowledgments

This study was supported by Inserm, CNRS, Féderation Recherche sur le Cerveau (FRC), Fondation Jérôme Lejeune (1728/2018 to BB and 1944/2019 to EL) and Agence Nationale de la Recherche (ANR-20-CE16-0016 to BB and EL and ANR-22-CE16-0011 to BB). SD was supported by Fédération Française pour la Recherche sur l’Epilepsie (FFRE). AT and TM were supported by Fraxa Research Foundation. S.L-G acknowledges the University’s Electron Microscopy facility (Centre Commun de Microscopie Appliquée, Université Côte d’Azur) and MICA Imaging platform Côte d’Azur supported by Université Côte d’Azur, Conseil Régional Sud Est PACA, Conseil Départemental and Gis Ibisa.

The authors are grateful to M. Capovilla for critical reading of the manuscript, A. Thomazeau, C. Gwizdek and F. Naro for discussion. They are indebted to F. Aguila for graphical artwork and to C. Boscagli for technical help.

