## Supplementary Files for "Defects in AMPAR trafficking and microglia activation underlie socio-cognitive deficits associated to decreased expression of Phosphodiesterase 2A"

**Behavioral experiments**

Experimental testing was performed between 8:00 and 12:00 each day during the 12-h light period

**Isolation-Induced Ultrasonic Vocalizations Test**

The test was performed as previously described (1). Briefly, each pup at PND10 was individually removed from the nest and placed into a Petri dish, located inside a sound-attenuating and temperature-controlled chamber. Pup ultrasonic vocalizations (USVs) were detected for 3 min by an ultrasound microphone (Avisoft Bioacoustics, Germany) sensitive to frequencies between 10 and 250 kHz and fixed at 10 cm above the arena. The emission of USVs was analyzed using the Avisoft Recorder software (Version 5.1).

**Homing Behavior Test**

The test was performed as previously described (1). At PND 13, the litter was separated from the dam and kept for 30 min in a temperature-controlled holding cage. Then, each mouse pup was placed into a Plexiglas box whose floor was covered for 1/3 with bedding from the pup's home cage and for 2/3 with clean bedding. The pup was located at the side of the box covered by clean bedding, and its behavior was video recorded for 5 min for subsequent analysis. The following parameters were scored: latency (s) to reach the home-cage bedding area; total time (s) spent by the pup in the nest bedding area.

**Social Interaction Test**

The test was performed as previously described (1). 28–30-day-old mice were individually habituated to the experimental apparatus (a Plexiglas cage measuring 30 × 30 × 30 cm) for 5 min the day before testing. On the test day, the animals were isolated for 2 h before testing, to enhance their social motivation and thus facilitate the expression of social interaction during testing. The test consisted in placing the experimental animal together with an unknown WT mouse of same age into the test cage for 10 min. The behavior of the mouse was recorded using an over-head camera and analyzed by a trained observer who was unaware of genotype using home-made counting software. The average frequency of total social activities (sniffing, following, mutual circle, grooming, crawling under/over), quantified as number of events during the 10 min testing session, was graphed.

**Elevated Zero Maze**

This test is used for assessing anxiety-related behaviors. The elevated circular maze has two enclosed areas opposite each other and two open areas. Before the test, mice were never manipulated to avoid habituation of height. At the beginning of the test, two month-old mice were gently invited to enter one of the closed areas and allowed to explore the maze for five minutes. Mouse behaviors were monitored via an over-head camera and analyzed for the time spent in the open areas and latency to first exit the initial closed area.

**Light-Dark Test**

The Light-Dark box consists of two chambers, one is dark and closed and the other one is brightly illuminated. At the start of the test, two-month-old mice were gently put in the dark chamber and were allowed to move freely between the two chambers. Mice behavior was monitored during five minutes *via* an over-head camera and analyzed for the time spent in the light chamber and latency to first exit the dark chamber.

**Novel Object Recognition**

Adult mice were habituated to the apparatus (20 x 38 cm) for ten minutes for two days before the test. On the third day, mice were placed individually into the arena with two identic objects (green bottle or coverslip box) for ten minutes during the learning phase. After this phase, the objects and the arena were cleaned with 70% ethanol and water. Five minutes or one hour after the learning phase, during the test phase, mice were free to explore a second time the same arena but with one of the objects replaced by another one from the other set. Object places and choice of set were randomly chosen to avoid preferences.

During these two phases the exploration time was manually assessed. Object exploration is defined when the mouse nose is closely directed toward the object, and not when animals are upon the object. Discrimination index is calculated during the test phase, corrected by the total time of exploration, as follows: (exploration time of new object – exploration time of old object)/(total time exploration). Animals having abnormal exploration time (less than ten seconds) were excluded from analysis.

**Morris Water Maze**

The apparatus used here was a circular pool of 90 cm of diameter filled with opaque water (23-24°C). A 8 cm diameter platform was used, which was visible during the Cue Task, hidden 1 cm under the water surface during the learning phase, and removed on Probe Test day. Four spatial cues for distal spatial navigation were present on the walls at the four cardinal points. For the Cue Task a visual cue was placed on top of the hidden platform to verify our animals' visual and motor capacity. Each day, mice were individually released into the pool four times, at each cardinal point, except for the Probe Test where they were tested one time from the West point. Mice had 90 seconds to find the platform, and had to stay for 30 seconds. If an animal did not find the platform, it was helped to reach it and had to stay there for 30 seconds.

The test was divided in two days of Cue Task, four days of learning the place of the hidden platform and one day of Probe Test. The latency to find and stay on the platform was measured for each animal, as well as the time spent into each quadrant for the Probe Test using the Anymaze software (AnyMaze, Stoelting, U.S.A.).

**Actimeter**

Mice were put individually into an actimeter cage (21 cm length × 11 cm width × 18 cm height) (Actimeter system, Imetronic, France) for 72 hours with food and water *ad libitum*. The actimeter cage consists of two laser beam rails, one above counting animal rearings and one below counting their general activity across the cage.

**Rotarod**

Motor performance was tested using an accelerating rotarod. Mice were tested using the acceleration mode (5–40 rpm) for three consecutive days with four trials per day. Mice were placed on the rod and the time that the individual mouse took to fall from the rod was measured.

**Electrophysiology**

The electrophysiology study was conducted by E-Phy-Science (https://www.e-phy-science.com/en/). Sagittal hippocampal brain slices were obtained using standard brain slicing methods. Mice were anesthetized with isoflurane and then decapitated. Brains were dissected out of the cranium and immediately immersed in ice-cold freshly prepared artificial CerebroSpinal Fluid (aCSF) containing: 117 mM NaCl, 4.7 mM KCl, 1.2 mM $MgCl_2$, 1.2 mM $NaH_2PO_4$, 2.5 mM $CaCl_2$, 25 mM $NaHCO_3$, and 11 mM D-glucose continuously oxygenated (pH = 7.4). Acute slices (350 µm thick) were prepared using a vibratome (VT 1000S; Leica Microsystems, Bannockburn, IL). Sections were incubated in standard aCSF (117 mM NaCl, 4.7 mM KCl, 1.2 mM $MgCl_2$, 1.2 mM $NaH_2PO_4$, 2.5 mM $CaCl_2$, 25 mM $NaHCO_3$, and 11 mM D-glucose) at room temperature for at least 1 hour before recordings. LTD measurements started between 90 and 120 minutes after excision of the brain.

For electrophysiological recordings, a single slice was placed in the recording chamber (room temperature), submerged and continuously superfused with gassed (95% $O_2$, 5% $CO_2$) aCSF at a constant rate (2 mL min−1) for the reminder of the experiment. aCSF was supplemented with 50 µM 2-amino-5-phosphonovaleric acid (AP5). Extracellular fEPSPs were recorded in the CA1 region of the hippocampus using a glass micropipette filled with aCSF. fEPSPs were evoked by the electrical stimulation of CA1 at 0.25 Hz with a glass stimulating electrode placed in the CA1 (100 µsec duration). Stable baseline fEPSPs were recorded by stimulating at 50% maximal field amplitude for a minimum of 10 min prior to beginning experiments (single stimulation every 10 s (0.1 Hz)). The same intensity of stimulation was kept for the reminder of the experiment. After a 10 min stable baseline period, LTD was induced by [S]-3,5-Dihydroxyphenylglycine [DHPG] bath application (100 µM, 5 minutes). The amount of LTD induced by metabotropic group I glutamate receptor (mGluR) was calculated 40 min after LTD induction by DHPG application. Signals were amplified with an Axopatch 200B amplifier (Molecular Devices, Union City, CA) digitized by a Digidata 1322A interface (Axon Instruments, Molecular Devices, US) and sampled at 10 kHz. Recordings were acquired using Clampex (Molecular Devices) and analyzed with Clampfit (Molecular Devices). Experimenters were blinded to treatment for all experiments.

**Untargeted metabolomics**

Twenty milligrams of snap-frozen tissue were crushed to a powder and solubilized in 700 µL of a methanol/water mixture (3:1, v/v) containing 50 µM 2-Morpholinoethanesulfonic acid (FC) as an internal standard (Cat. # 341-01622; Dojindo, Tokyo, Japan). The samples were homogenized using 0.5g of glass beads (Retsch, Cat. # 22.222.0002; Retsch, Haan, Germany) and vortexed for 1 minute. They were further processed using the Tissue Homogenizer Precellus 24 (Brevet Bertin Technologies, Aix-en-Provence, France) for 30 seconds. The homogenate was then centrifuged at 15,000*g*/4°C for 15 minutes, and the supernatant was loaded onto a Captiva EMR plate (Agilent, Cat.# 5190-1001; Santa Clara, CA, USA), assembled on the Vacuum Manifold (Agilent, Cat.# A796), together with the Deep Well collection plate (Agilent, Cat.# A696001000). The flow-through was collected and dried using lyophilization.

For metabolomic analysis, the samples were diluted in 200 µL of water, and 1 µL was injected into an LC-MS system consisting of a UHPLC (Vanquish Flex; Thermo, Bremen, Germany) coupled to an orbitrap MS (QExactive Plus, Thermo). The analytical column used was a ZORBAX-Pursuit 3 PFP column (Agilent; Cat. No.: A3051150X020) with dimensions 150 x 2.0 mm. The mobile phases were composed as follows: A: 99.9% water, 0.1% formic acid (Sigma Aldrich, Saint Louis, MO, USA; Cat.# 33015); B: 99.9% acetonitrile (Biosolve BV, Valkenswaard, Niederlande; Cat.# 001204102BS), 0.1 % formic acid. The gradient was as follows: 0 min (0 % B), 2 min (25 % B), 11 min (35 % B), 15 min (95 % B), 20 min (95 % B), 21 min (0 % B). The flow rate was set at 0.25 ml/min, and the column temperature was maintained at 40 °C. The UHPLC was connected to the MS via an ESI ion source with the following parameters in positive mode: sheath gas flow 40, auxiliary gas flow 10, spray voltage 3.8 kV, capillary temperature 350 °C, S-lens RF level 55, and auxiliary gas heater 200 °C. The MS acquisition method followed a data-dependent approach, where the top 5 MS ions with a default charge of 1 were selected for MS/MS fragmentation. The acquisition parameters were set as follows: scan range 80 to 750 m/z, resolution 17,500, maximum IT 50 ms, AGC target 1e5, isolation window 4 m/z, collision energy stepped at 20, 40, and 60.

The raw data were imported into Compound Discoverer (CD) software (Thermo, Version 3.3), where they underwent processing using the default untargeted metabolomics workflow. Statistical analysis of modulated metabolites was performed using the CD software, with a fold-change threshold of 2 and a p-value <0.05 being considered as significant.

**Supp. Fig. 1 : Complementary behavioral study of male *Pde2a*$^{+/-}$ mice.**

**(A,B)** WT and *Pde2a*$^{+/-}$ mice have the same weight at P10 and P13 (n=14-15). **c** *Pde2a*$^{+/-}$ mice do not show anxiety-like behavior in Light Dark test, as latency of first exit from the dark box and time spent in the light box are equivalent between the two groups (n=11-13). **(D,E)** Exploration time during the learning phase of NOR test are not significantly different between WT and *Pde2a*$^{+/-}$ mice (n=9-14). **(F)** *Pde2a*$^{+/-}$ mice show significant longer time to reach the visible platform the first day, but show no difference in learning the hidden platform location in Morris Water Maze after 4 days of training compared to WT (n=7-11). **(G)** Total activity and day/night cycle in 3 days are not altered in *Pde2a*$^{+/-}$ mice **(H)** but they tend to do more rearing during this time (n=10-14). **(I)** WT and *Pde2a*$^{+/-}$ mice have the same latency to fall during the three days of rotarod test (n=8-9). The data are represented as means ± SEM and analyzed using the Mann-Whitney test and a Two-Way ANOVA with Tukey's multiple comparisons test **(G,H)** (adjusted p value: *P<0.05, **P<0.01, ***P<0.001).

**Supp. Fig. 2 : Female *Pde2a*$^{+/-}$ mice present milder behavior phenotypes compared to males.**

**(A,B)** Female *Pde2a*$^{+/-}$ mice do not show deficits in USV and Homing test, respectively (n=13-17). **(C)** At one-month, female *Pde2a*$^{+/-}$ interacts significantly but slightly less with their counterparts compared to WT (n=10-11), **(D)** but at four months this difference is no more visible (n=8-10). **(E)** Female *Pde2a*$^{+/-}$ mice don't show anxiety-like behavior in EZM, as latency of first exit and time spent in open arm are equivalent to WT mice (n=9). **(F)** Female *Pde2a*$^{+/-}$ mice have no preference for novel object in NOR test with 5 min interval, with discrimination index near zero (n=7-9), while exploration time in **(G)** learning and **(H)** test phase are equivalent to WT mice.

**Supp. Fig. 3 : Microglia in *Pde2a*$^{+/-}$ brains.**

**(A)** Representative image of WT and *Pde2a*$^{+/-}$ cortex immuno-stained with DAPI (blue) and IBA1 (green) (bar scale: 200µm). **(B)** Western Blot and quantification show a significant increase IBA1 in *Pde2a*$^{+/-}$ cortex at two-weeks-old **(C)** but not at one month of age compared to WT (n=4-6). **(D)** In confocal image to analyze microglia morphology, *Pde2a*$^{+/-}$ cortex contains more IBA1 positive cells (n=5-6). **(E,F)** Relative mRNA expression of various inflammatory markers in hippocampus and cortex of WT and *Pde2a*$^{+/-}$ mice at one month of age (n=6-9). CCL12 expression could not be detected in hippocampus at this age. The data are represented as means ± SEM and analyzed using unpaired t test. Mann-Whitney test was used to compare groups that did not pass test for normality. (p value : *P<0.05, **P<0.01)


1. Maurin T, Melancia F, Jarjat M, Castro L, Costa L, Delhaye S, et al. Involvement of Phosphodiesterase 2A Activity in the Pathophysiology of Fragile X Syndrome. Cereb Cortex. 2019 Jul 22;29(8):3241–52.

Supplementary Figure 1

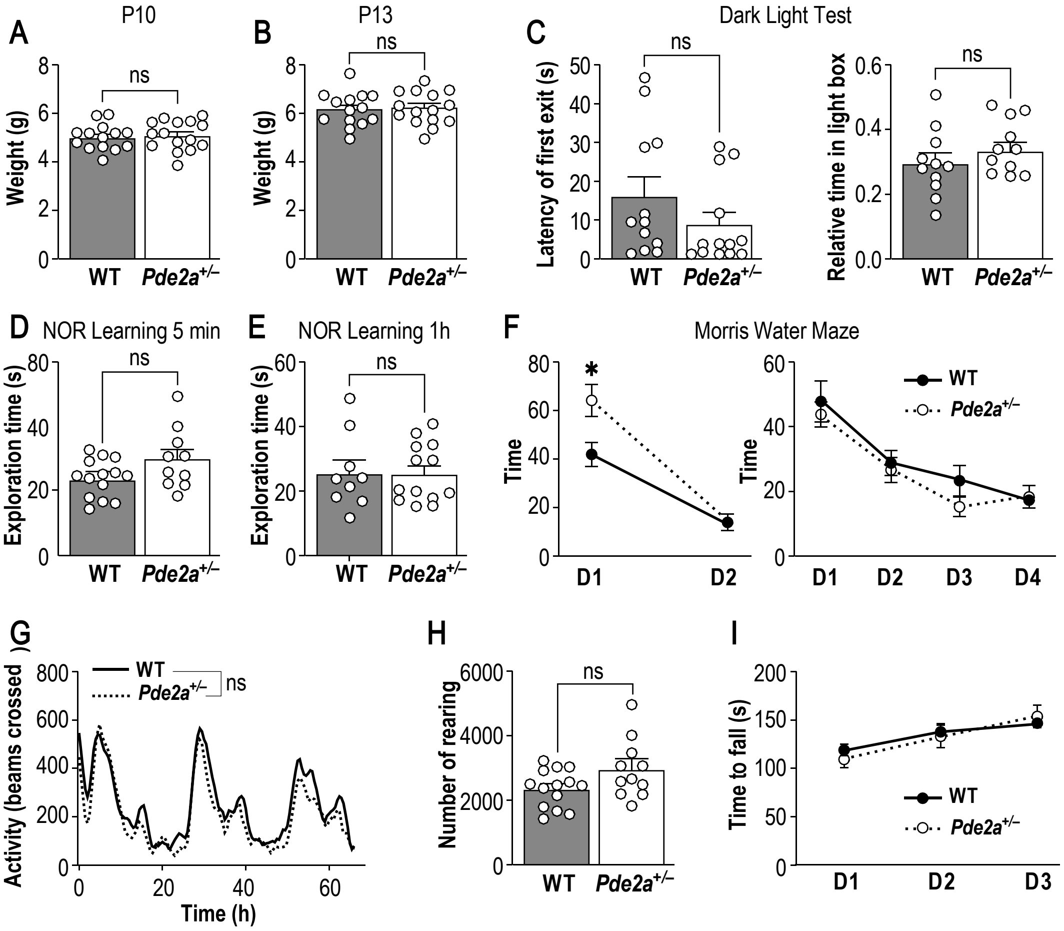

Supplementary Figure 2

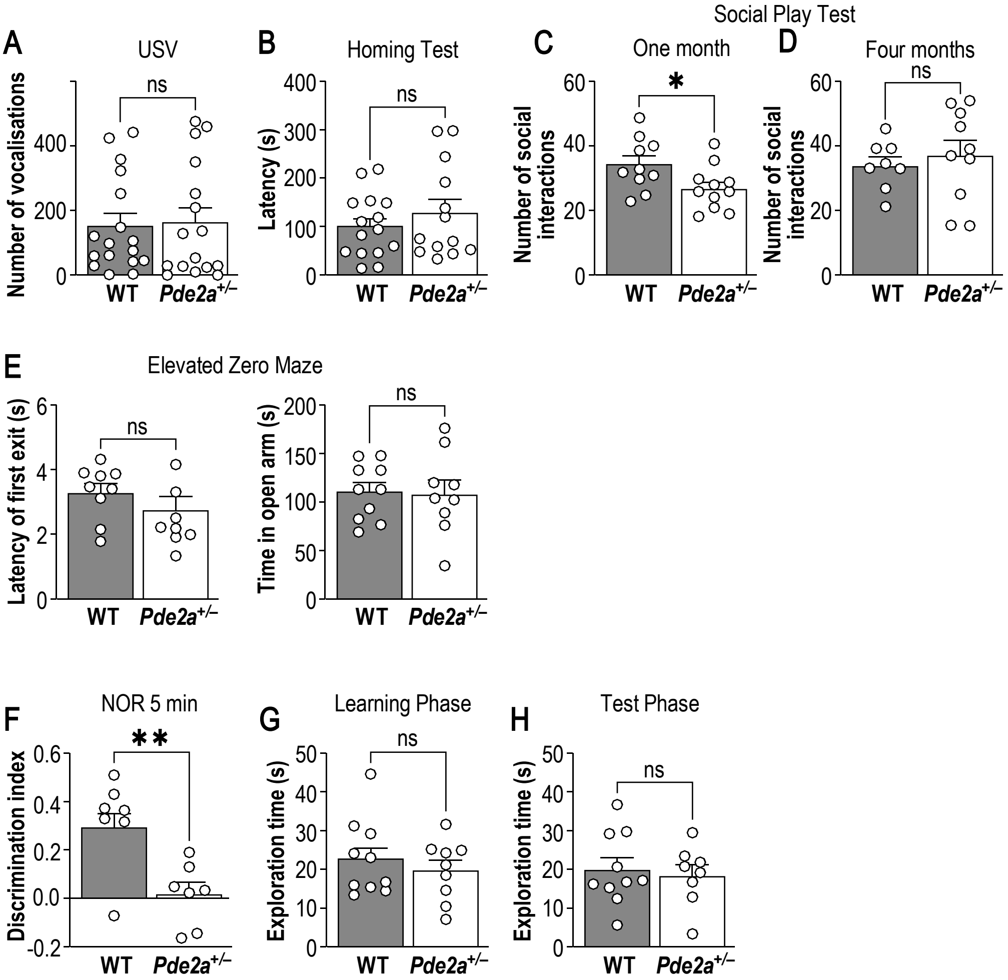

### Supplementary Figure 3

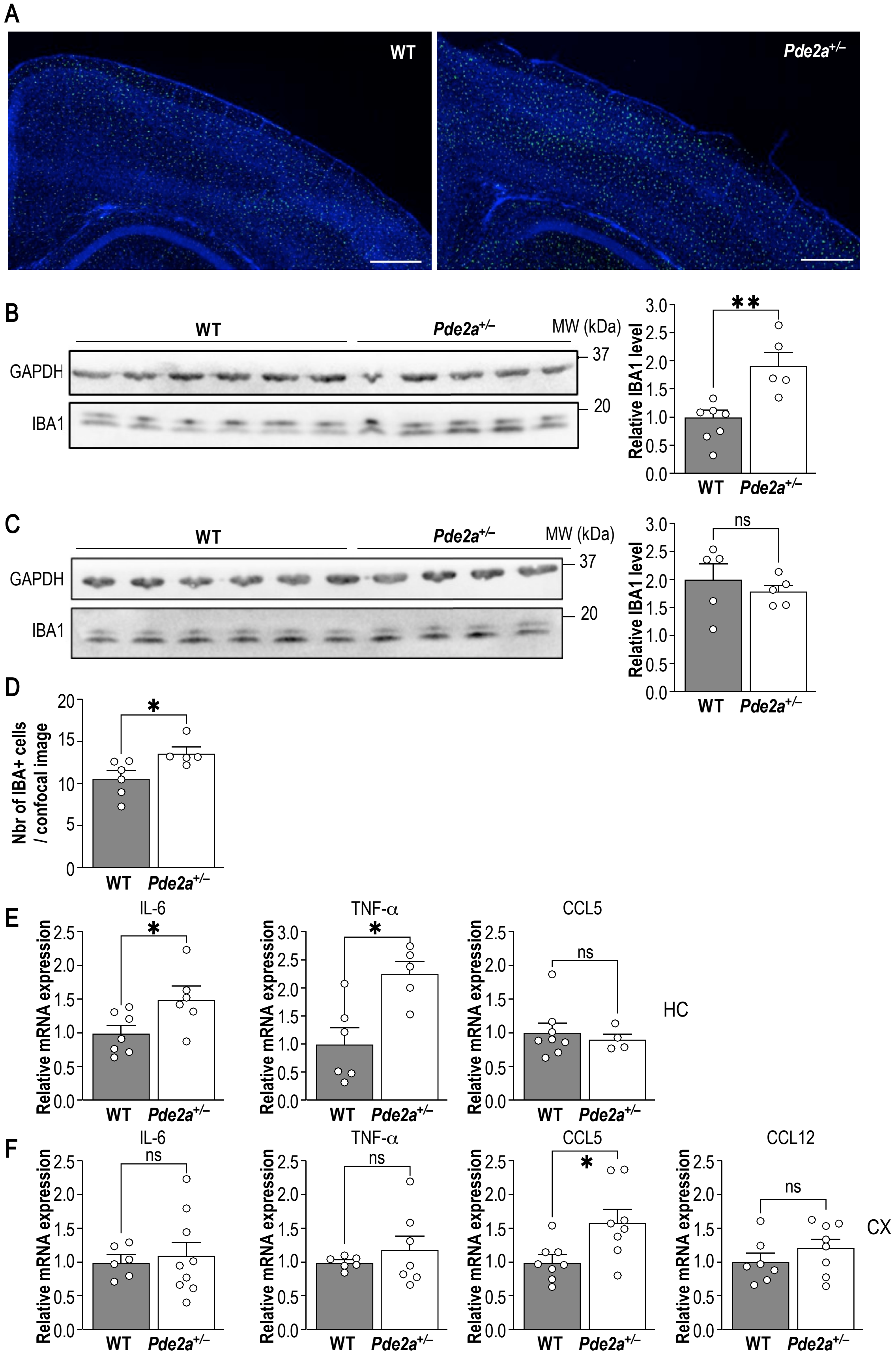

| **CORTEX $p<0.05$** | **Cortex *Pde2a*$^{+/-}$ (>2-fold)** |
|---|---|
| Ggamma-L-Glutamyl-L-glutamic acid | upregulated |
| L-Pyroglutamic acid | upregulated |
| (3R)-beta-Leucine | upregulated |
| L-Glutamic acid | upregulated |
| N-Acetylaspartylglutamic acid | upregulated |
| Allopurinol | upregulated |
| Inosine | upregulated |
| Tropinone | upregulated |
| Thymine | upregulated |
| 3-Dehydroxycarnitine | upregulated |
| 2'_4'-Dihydroxyacetophenone | upregulated |
| L-Tryptophan | upregulated |
| N-Acetyl-L-phenylalanine | upregulated |
| Glutathione | upregulated |
| IMP | upregulated |

**Supp. Table 1.** List of metabolites significantly upregulated (more than 2-fold) in cortex extracts from *Pde2a*$^{+/-}$ mice compared to WT.

| | Total | Expected | Hits | Raw p | Holm p | FDR |
|---|---|---|---|---|---|---|
| Glutathione Metabolism | 21 | 0.31 | 3 | 2.89E-03 | 2.83E-01 | 2.83E-01 |
| Purine Metabolism | 74 | 1.08 | 3 | 8.79E-02 | 1.00E+00 | 1.00E+00 |
| Aspartate Metabolism | 35 | 0.51 | 2 | 9.02E-02 | 1.00E+00 | 1.00E+00 |
| Malate-Aspartate Shuttle | 10 | 0.15 | 1 | 1.38E-01 | 1.00E+00 | 1.00E+00 |
| Pyruvaldehyde Degradation | 10 | 0.15 | 1 | 1.38E-01 | 1.00E+00 | 1.00E+00 |
| Glutamate Metabolism | 49 | 0.72 | 2 | 1.59E-01 | 1.00E+00 | 1.00E+00 |
| Glucose-Alanine Cycle | 13 | 0.19 | 1 | 1.76E-01 | 1.00E+00 | 1.00E+00 |
| Tryptophan Metabolism | 60 | 0.88 | 2 | 2.18E-01 | 1.00E+00 | 1.00E+00 |
| Alanine Metabolism | 17 | 0.25 | 1 | 2.23E-01 | 1.00E+00 | 1.00E+00 |
| Arachidonic Acid Metabolism | 69 | 1.01 | 2 | 2.68E-01 | 1.00E+00 | 1.00E+00 |
| Cysteine Metabolism | 26 | 0.38 | 1 | 3.22E-01 | 1.00E+00 | 1.00E+00 |
| Phenylalanine and Tyrosine Metabolism | 28 | 0.41 | 1 | 3.42E-01 | 1.00E+00 | 1.00E+00 |
| Folate Metabolism | 29 | 0.42 | 1 | 3.52E-01 | 1.00E+00 | 1.00E+00 |
| Urea Cycle | 29 | 0.42 | 1 | 3.52E-01 | 1.00E+00 | 1.00E+00 |
| Lysine Degradation | 30 | 0.44 | 1 | 3.62E-01 | 1.00E+00 | 1.00E+00 |
| Ammonia Recycling | 32 | 0.47 | 1 | 3.81E-01 | 1.00E+00 | 1.00E+00 |
| Amino Sugar Metabolism | 33 | 0.48 | 1 | 3.90E-01 | 1.00E+00 | 1.00E+00 |
| Beta-Alanine Metabolism | 34 | 0.50 | 1 | 4.00E-01 | 1.00E+00 | 1.00E+00 |
| Nicotinate and Nicotinamide Metabolism | 37 | 0.54 | 1 | 4.26E-01 | 1.00E+00 | 1.00E+00 |
| Propanoate Metabolism | 42 | 0.61 | 1 | 4.69E-01 | 1.00E+00 | 1.00E+00 |
| Histidine Metabolism | 43 | 0.63 | 1 | 4.77E-01 | 1.00E+00 | 1.00E+00 |
| Pyruvate Metabolism | 48 | 0.70 | 1 | 5.16E-01 | 1.00E+00 | 1.00E+00 |
| Arginine and Proline Metabolism | 53 | 0.78 | 1 | 5.52E-01 | 1.00E+00 | 1.00E+00 |
| Warburg Effect | 58 | 0.85 | 1 | 5.86E-01 | 1.00E+00 | 1.00E+00 |
| Glycine and Serine Metabolism | 59 | 0.86 | 1 | 5.92E-01 | 1.00E+00 | 1.00E+00 |
| Pyrimidine Metabolism | 59 | 0.86 | 1 | 5.92E-01 | 1.00E+00 | 1.00E+00 |
| Valine, Leucine and Isoleucine Degradation | 60 | 0.88 | 1 | 5.98E-01 | 1.00E+00 | 1.00E+00 |
| Tyrosine Metabolism | 72 | 1.05 | 1 | 6.68E-01 | 1.00E+00 | 1.00E+00 |

**Supp. Table 2.** Results from Over Representation Analysis of differentially abundant metabolites.

| | |
|---|---|
| PDE2A F1 | CTGCCTGATGGTGAAGAAAGGCTA |
| PDE2A F2 | GGGCCAGCTCATTCCTCCCACTCAT |
| PDE2A R | TGAGCAGACCCCTTATGGAAGGTG |
| IL-6 F | CCCAATTTCCAATGCTCTCCT |
| IL-6 R | GAATTGGATGGTCTTGGTCC |
| TNF -α F | GGTGACCAGGCTGTCGCTAC |
| TNF -α R | AGGGCAATTACAGTCACGGC |
| CCL5 F | ACACCACTCCCTGCTGCTTT |
| CCL5 R | AAATACTCCTTGACGTGGGCA |
| CCL12 F | ACACTGGTTCCTGACTCCTCT |
| CCL12 R | ACCTGAGGACTGATGGTGGT |
| TBP F | AGGCCAGACCCCACAACTC |
| TBP R | GGGTGGTGCCTGGCAA |

**Supp. Table 3**. Oligos for genotyping PCR and qPCR.
